# EpipwR: Efficient Power Analysis for EWAS with Continuous Outcomes

**DOI:** 10.1101/2024.09.06.611713

**Authors:** Jackson Barth, Austin W. Reynolds

**Author notes:** Corresponding author. Jackson.

## Abstract

**Motivation:** Epigenome-wide association studies (EWAS) have emerged as a popular way to investigate the pathophysiology of complex diseases and to assist in bridging the gap between genotypes and phenotypes. Despite the increasing popularity of EWAS, very few tools exist to aid researchers in power estimation and those are limited to case-control studies. The existence of user-friendly tools, expanding power calculation functionality to additional study designs would be a significant aid to researchers planning EWAS.

**Results:** We introduce EpipwR, an open-source R package that can efficiently estimate power for EWAS with continuous outcomes. EpipwR uses a quasi-simulated approach, meaning that data is generated only for CpG sites with methylation associated with the outcome, while p-values are generated directly for those with no association (when necessary). Like existing EWAS power calculators, reference datasets of empirical EWAS are used to guide the data generation process. Two simulation studies show the effect of the selected empirical dataset on the generated correlations and the relative speed of EpipwR compared to similar approaches.

**Availability and Implementation:** The EpipwR R-package is currently available for download at github.com/jbarth216/EpipwR.

## 1. Introduction

Over the past decade, epigenome-wide association studies (EWAS) have emerged as the dominant way for researchers to investigate the relationship between epigenetic markers and phenotypes at the genome-wide level. DNA methylation has become the most widely studied epigenetic mechanism because of its ease to assay through existing microarray and sequencing technology, its ubiquity across the genome, and its impact on gene expression. DNA methylation, the process whereby a methyl group is added to a nucleotide changing its availability for transcription, occurs most often in the context of cytosine-phosphate-guanine (CpG) dinucleotides in millions of locations around the genome. To assay this variation at a population scale, most EWAS use microarray-based platforms for assessing DNAm, such as the Illumina Infinium HumanMethylation450 BeadChip (covering ∼450,000 CpG sites in the human genome) or the Illumina Infinium MethylationEPIC Array (covering *>*850,000 CpGs). Both arrays assess methylation at single-CpG resolution, quantified using a methylation *β*-value; an approximately continuously-distributed measure that reflects the level of methylation at a specific locus, ranging from 0 (unmethylated) to 1 (methylated) (Bibikova et al., 2006). The development of these arrays has precipitated a huge interest in studying DNA methylation in the context of human health and disease (Murphy and Mill, 2014; Rakyan et al., 2011; Wei et al., 2021). The need for power calculations to inform EWAS study design has often been reiterated since their advent (Mansell et al., 2019; Michels et al., 2013; Tsai and Bell, 2015), but few tools have been created to aid researchers.

It is now widely accepted that formal power calculation and sample size justification is an essential part of genomic study design to ensure meaningful findings are reported. This has motivated the development of numerous statistical power evaluation tools for genome- and transcriptome-wide association studies, including: the GAS Power Calculator (Johnson and Abecasis, 2017), GWAPower (Feng et al., 2011), RnaSeqSampleSize (Zhao et al., 2018), and RNAseqPS (Guo et al., 2014). However, surprisingly few tools for EWAS power evaluation have been developed, despite substantial work in the areas of DNA methylation data QC, normalization, and analysis (Aryee et al., 2014; Chakravarthy et al., 2018; Fortin et al., 2014; Houseman et al., 2012; Jaffe et al., 2012; Morris et al., 2014; Peters et al., 2015; Song and Kuan, 2022; Teschendorff et al., 2013, 2017). The pwrEWAS package (Graw et al., 2019) is perhaps the only tool currently available to researchers for easily estimating power under a variety of conditions (e.g. sample size, effect size, false discovery rate threshold). Even with the availability of this tool, most EWAS continue to be conducted without formal power analyses, resulting in potentially under- and over-powered studies (Tsai and Bell, 2015). Furthermore, pwrEWAS is currently limited to case-control study designs, limiting the ability of researchers to easily estimate sample size requirements during study design.

The scope of EWAS, however, have expanded beyond two-group comparison designs into the realm of complex traits. For example, the relationship between age and CpG methylation has been extensively studied over the past decade (Hannum et al., 2013; Horvath, 2013; Horvath and Raj, 2018), with many applications to other health outcomes (Belsky et al., 2020, 2022; Levine et al., 2018; Lu et al., 2019a,b, 2022). More recently, EWAS have investigated CpG methylation correlated with a number of quantitative traits; including blood metabolites (Gomez-Alonso et al., 2021; Jhun et al., 2021), body mass (Aslibekyan et al., 2015; Demerath et al., 2015; Wahl et al., 2017), lung function (Herrera-Luis et al., 2022), and even behavioral health (Barbu et al., 2022; Montalvo-Ortiz et al., 2022; Raffington et al., 2023). As the number and complexity of phenotypes being studied with EWAS continues to grow, so too does the research community’s need for appropriate and easily accessible power calculation tools.

Here we introduce EpipwR, a publicly available tool for comprehensive EWAS power evaluation in the context of continuous outcome variables. EpipwR is a quasi-simulated approach to power analysis, in that data based on empirical EWAS are generated only for non-null distributions while false positive p-values are generated directly using classic statistical theory. In this way, EpipwR leverages the unique form of EWAS data while running much faster than a fully-simulated method. EpipwR was written using the R statistical programming language (R Core Team, 2021) and the package is available at https://github.com/jbarth216/EpipwR.

## 2. Methods

Suppose an EWAS has *K* CpG sites and *n* distinct patient samples. Let *β*_*ik*_ be the proportion of DNA methylation from patient *i*, CpG *k*, where it is assumed that *β*_*ik*_ from a common CpG site are independently and identically distributed from some beta distribution, *β_·k_ ∼ Beta*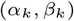. Of interest to researchers is a continuous phenotype *Y*_*i*_ which is assumed to be correlated with the DNA methylation of some CpG sites, *ρ*_*k*_ = *cor*(*Y, M*_*·k*_), where *M*_*ik*_ = *logit*(*β*_*ik*_). While *M*_*ik*_ is usually closer to normality than *β*_*ik*_ (Du et al., 2010), it should be noted that skewed distributions of *β*_*ik*_ can still yield non-normal *M*_*ik*_. Finally, *K* can be split into true nulls (*K*_*n*_ of *K* have *ρ*_*k*_ = 0) and false nulls (*K*_*m*_ of *K* have *ρ*_*k*_ 0). The goal of EpipwR is to evaluate the power of this analysis, defined as the proportion of false null hypotheses among *K*_*m*_ that are correctly identified. The remainder of this section outlines and justifies the methodology implemented in EpipwR.

### 2.1. Data Generation

The first step in calculating power for the scenario described above is to generate methylation data, *β*_*ik*_. In an effort to maintain consitency across tools used in epigenomics, EpipwR applies the same method outlined by Graw et al. (2019) in pwrEWAS to generate the methylated proportions. To determine plausible distributions for each non-null CpG site (which will then be used to generate *β*_*ik*_), EpipwR relies on empirical EWAS reference data. One of twelve publicly available EWAS of different tissue types is selected by the user based on relevance to the study being planned. Each dataset corresponds to a different list of beta distribution parameters 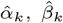 estimated through method of moments on every CpG site in the selected dataset. For every truly correlated CpG site in the planned study, EpipwR samples *K*_*m*_ pairs of these parameters with replacement to be used to generate the methylation data. Unlike pwrEWAS which is a fully simulated approach, methylation datasets are not generated for the true nulls in EpipwR (see section 2.2), which significantly reduces computation time and memory load.

Once the beta distributions have been identified, the next step is to generate *n* samples of (*Y, β*_*·*1_, *β*_*·*2_, …, *β*_.*K_m_*_). EpipwR assumes that all *β*_*ik*_ are conditionally independent on *Y* such that *Cov*(*β*_*·k*_, *β*_*·l*_|*Y*) = 0 for the *k*^*th*^ and *l*^*th*^ CpG sites. This is a common simplifying assumption in power analysis, and one that removes the responsibility from the researcher of identifying a valid covariance structure for potentially thousands of *β*_*·k*_. This assumption allows *Y* to be sampled marginally, and then the subsequent *β*_*ik*_|*Y* can be sampled concurrently. EpipwR also assumes that *Y* is normally distributed and assumes without loss of generality that it follows a standard normal distribution. Therefore, EpipwR initially generates *n* samples of *Y ∼ N* (0, 1). To conditionally sample *β*_*ik*_, EpipwR utilizes a version of rank correlation closely based on Ruscio and Kaczetow (2008). First, *n* samples of *X*_*k*_|*Y* are generated, where *X*_*k*_ has a marginal standard normal distribution with *ρ*_*k*_ = *cor*(*X*_*k*_, *Y*). The percentiles of each *X*_*ik*_ are then used to systematically generate *β*_*ik*_, such that 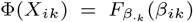where Φ is the standard normal cumulative distribution function (CDF) and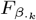 is the CDF of *β*_*ik*_ *∼ Beta* 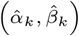 with parameters sampled from the empirical EWAS data. For example, suppose *x*_*ik*_ is generated to be *−*0.8416. Since this is the 20th percentile value of the standard normal distribution, then *β*_*ik*_ is assignedthe 20th percentile value of its corresponding beta distribution. The *β*_*ik*_ are then converted to *M*_*ik*_ via the logit transformation.

Taking the approach described in Ruscio and Kaczetow (2008) for generating non-normal correlated data would involve randomly generated beta samples, which are then reordered based on the ranks of *X*_*ik*_. EpipwR leverages the inverse CDF of the beta distribution to directly convert *X*_*ik*_ to *β*_*ik*_. While the computational resources of each approach is similar, using the inverse CDF tends to be slightly more accurate and has a standard error closer to what would be expected under normal conditions (see section 3.1). Additionally, the method in Ruscio and Kaczetow (2008) is an iterative approach, adjusting *ρ*_*k*_ and repeating the process until *r*_*k*_ (the sample correlation) reaches the desired level. EpipwR does not use an iterative approach but rather accepts the first generated correlation for each CpG. This is not neccesarily because EpipwR will always produce *ρ*_*k*_ = *r*_*k*_ (although it is more accurate on average) but because this is a more accurate reflection of reality. When employing Pearson correlation (or any parametric measure) to assess the association between two quantitative variables, it is essential to acknowledge the underlying assumption of linearity in their relationship. However, this assumption may not always hold true, particularly when one of the distributions exhibits skewness (as is the case with the reference data-see figure 1). In such instances, only a segment of the relationship may adhere to linearity, leaving the remaining portion inaccurately estimated by the test. Therefore, if *ρ*_*k*_ is meant to describe the entrie relationship between *Y* and the DNA methylation, the generation of a linear correlation coefficient *r*_*k*_ slightly lower than anticipated under normal conditions is not a flaw but rather a distinctive feature of the method. Simulation results in Section 3.1 shows that heightened skewness in *β*_*·k*_ corresponds with a more pronounced underestimation in *r*_*k*_ compared to perfectly normal conditions.

**Fig 1.**
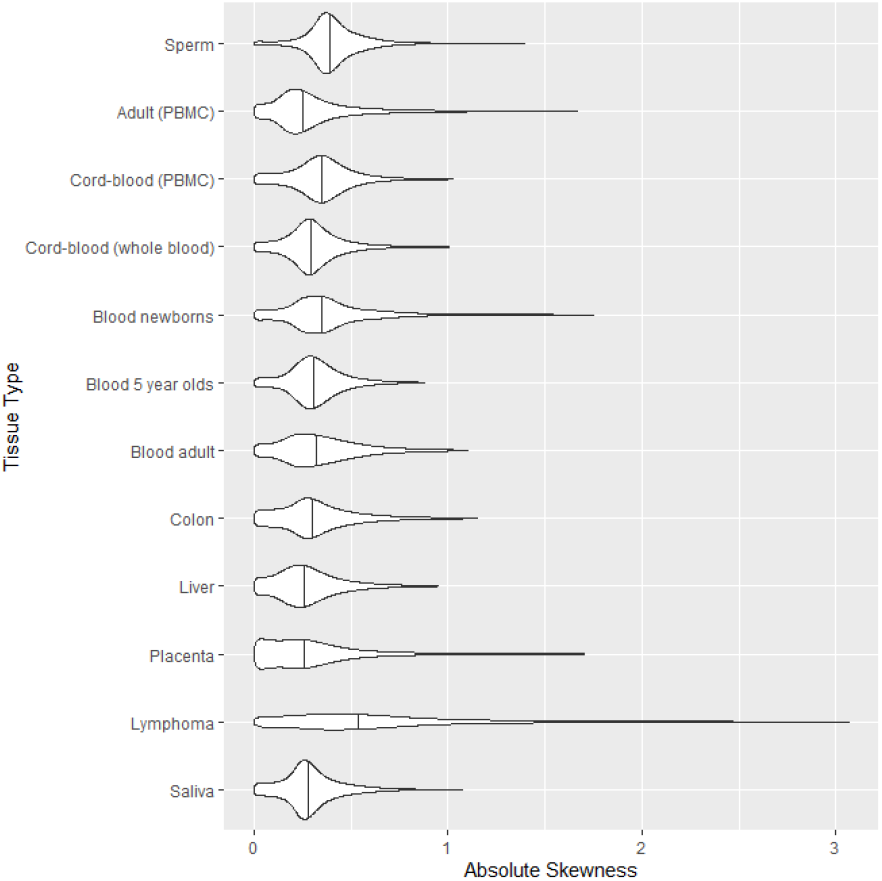
The violin plots show the distributions of absolute skewness across each of the 12 reference datasets used by EpipwR. For ease of viewing, the plots only show up to the 99th percentile of each dataset. The vertical lines display the median of each dataset

In setting the target correlation *ρ*_*k*_, researchers have the option of choosing a fixed correlation for all CpG sites such that *µ*_*ρ*_ = *ρ*_*k*_ or of identifying parameters of a truncated normal distribution to randomly generate the correlation of each associated CpG site:

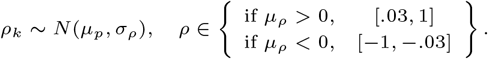

For the non-null tests, *ρ*_*k*_ cannot be in the range (*−*.03, .03), preserving practical significance.

### 2.2. Calculating p-values

For the *K*_*m*_ associated CpG sites, the p-value calculations are standard. The sample correlation for each CpG site, *r*_*k*_, is first calculated based on the chosen method (Pearson, Kendall or Spearman) for each dataset, and then p-values are similarly determined. For the non-parametric tests, exact p-values are calculated when the data have no ties (this is often the case) and are estimated with parametric methodology when ties are present. Becasue EpipwR (like most power analysis software) does not take into account dependence between CpG sites unrelated to *Y*, using EpipwR with the pearson test would yield the same results if the analysis was instead run using specialized software such as limma (Ritchie et al., 2015).

If a bonferonni adjustment is used to control for the family-wise Type I error rate, then p-values for the *K*_*n*_ null tests can be safely ignored. If instead users wish to control for a false discovery rate (FDR), then additional p-values for the non-significant associations must be considered. For the *K*_*n*_ CpG sites with a true null, computational resources are reduced significantly by generating p-values from a uniform distribution on the interval (0, 1), a well-established result in statistical theory. However, even this approach can become computationally taxing if *K*_*n*_ is large and the sampling is repeated on several datasets (*N >* 100). To further improve computational efficiency, EpipwR then calculates the largest order statistic from the *K*_*n*_ p-values with a probability of at least 1 *×* 10^*−*10^ of moving the cutoff and altering the power calculation (call this *ξ*). For each dataset, EpipwR then draws only the first *ξ* order statistics from *K*_*m*_ *U* (0, 1) samples, each of which can be sampled directly from a beta distribution (For a large number of tests, *xi* is around 30-40). All other p-values are assumed to be too large to alter the FDR cutoff.

### 2.3. Calculating Power

EpipwR uses a traditional calculation to evaluate power, where power is the percentage of the *K*_*m*_ truly associated CpG sites found to be significant. False positives are ignored in the power calculation (although if FDR is chosen, the false positives can effect the threshold). This calculation is repeated for each of the *N* datasets. The average across all *N* datasets is accepted as the average power.

Rather than specifying a specific number of datasets *N*, EpipwR uses a dynamic process to determine *N* as it runs. First, EpipwR calculates power for 20 different datasets (the mininum value for *N*). Beginning at the 20th dataset, it then calculates the margin of error of a 95% confidence interval for average power using the below formula,

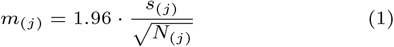

where *s*_(*j*)_ is the sample standard deviation of power through the first *N*_(*j*)_ generated datasets. If *m*_(*j*)_ is less than a user-specified margin of error (the default is .03), the process stops and uses the results from the *N*_(*j*)_ generated datasets to calculate average power. Otherwise, a new dataset is generated and the algorithm repeats, this time with *N*_(*j*+1)_ = *N*_(*j*)_ + 1 datasets. The process continues until *m*_(*j*)_ is small enough or if *N*_(*j*)_ reaches a user-specified maximum (default is 1,000) to limit the strain on computational resources.

### 2.4. EpipwR in Practice

There are two main functions in the EpipwR package, one that calculates power under various scenarios (EpipwR cont) and one that plots the results of the power analysis (EpipwR plot). EpipwR cont takes 12 inputs, as indicated in table 1. Since the sample size and *µ*_*ρ*_ inputs can take on multiple values, EpipwR calculates power separately for all scenarios under the cartesian product of these two vectors. The output of EpipwR cont is a dataframe with 1 row for every combination of n and rho mu, and columns tracking the average power, number of datasets used and the standard error of the average power. EpipwR plot takes this dataframe and plots 95% confidence intervals using ggplot2 (Wickham, 2016).

**Table 1.** EpipwR cont inputs.

| Name | Description | Notes |
| --- | --- | --- |
| dm | Amount of non-null tests |  |
| Total | Amount of all tests |  |
| n | Sample size(s) at which power is calculated | Accepts a vector of sample sizes |
| fdr_fwer | family-wise type I error or false discovery rate (FDR) | Depends on use_fdr |
| rho_mu | Average value of $\rho$ for non-null tests | Accepts a vector of averages |
| rho_sd = 0 | Standard deviation of $\rho$ for non-null tests | 0 implies that $\rho_k = \text{rho\_mu}$ for all non-null tests |
| Tissue = "Saliva" | Reference data used to generate $\beta_{ik}$ | See package for valid options |
| Nmax = 1000 | Maximum number of datasets generated | Minimum number is fixed at 20 |
| MOE = .03 | Target margin of error for a 95% confidence interval | See section 2.3 |
| test = "pearson" | Type of correlation/test utilized | Can be pearson, kendall or spearman |
| use_fdr = T | Indication that fdr_fwer should be treated as FDR | If false, fdr_fwer is treated as FWER |
| Suppress_updates = F | If T, removes intermediate status updates | Has no impact on the results |

Figure 2 demonstrates the EpipwR workflow. In this scenario, researchers plan an EWAS with 100,000 CpG sites, of which 500 are expected to have non-null correlations with some phenotype of interest. The researchers wish to test sample sizes of 100-200 at denominations of 25 and *µ*_*ρ*_ *∈* (0.3, 0.35, 0.4) where *ρ*_*k*_ are fixed at these values. At a 5% false discovery rate, the plots on the right-hand side of figure 2 show that sample sizes of 125 and 150 are high enough to acheive 80% and 90% power (respectively) if *µ*_*ρ*_ = 0.4, whereas larger sample sizes are needed for smaller correlations.

**Fig 2.**
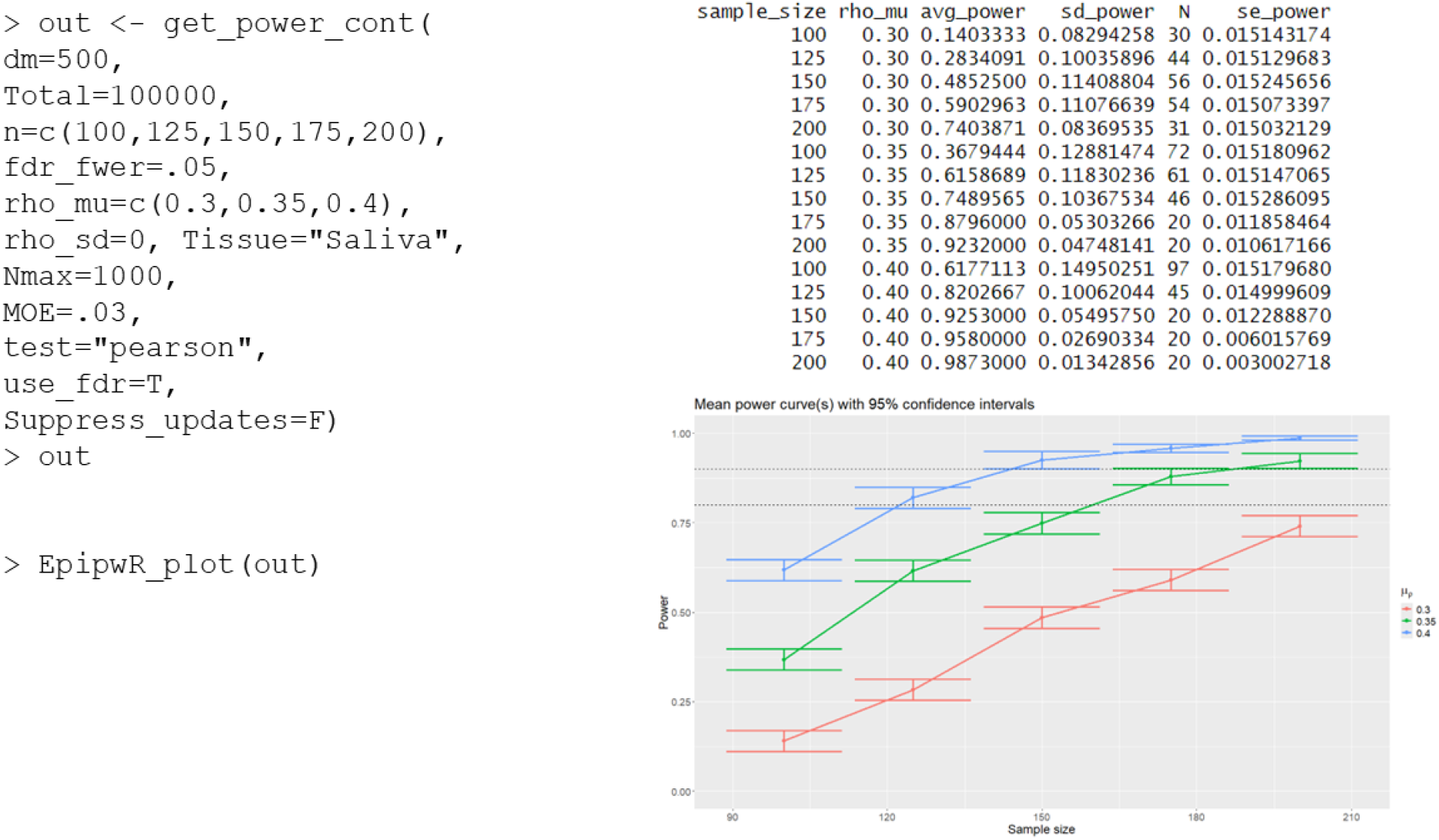
Code and output of the primary EpipwR functions. The get power cont() function produces the R-console table on the right-hand side, while EpipwR plot() produces the error bar plot

## 3. Results

### 3.1. Generation of correlated datasets

To evaluate the efficacy of our method to produce dependent data, a small simulation was conducted to observe the large-sample behavior of the sample correlations under differing beta distributions. Sample datasets for CpG methylation were generated based on 800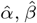; estimates from the saliva dataset, drawn using stratified sampling based on the estimated skewness (*γ*) of the distribution. Specifically, 100 pairs of estimates were drawn such that |*γ*| *∈* (0, 0.01), another 100 were drawn such that |*γ*| *∈* (0.01, 0.5) and so on for a variety of ranges (see the *x*-axis of figure 3). Correlated data were then generated based on sample sizes *n ∈* (10, 30, 50, 100, 150, 200) and “true” correlation *ρ* from 0.1 to 0.9 in increments of 0.1. For each simulation setting, 1,000 correlated datasets were generated based on the method described in section 2.1. The average and standard deviation of the sample correlations were then stored.

**Fig 3.**
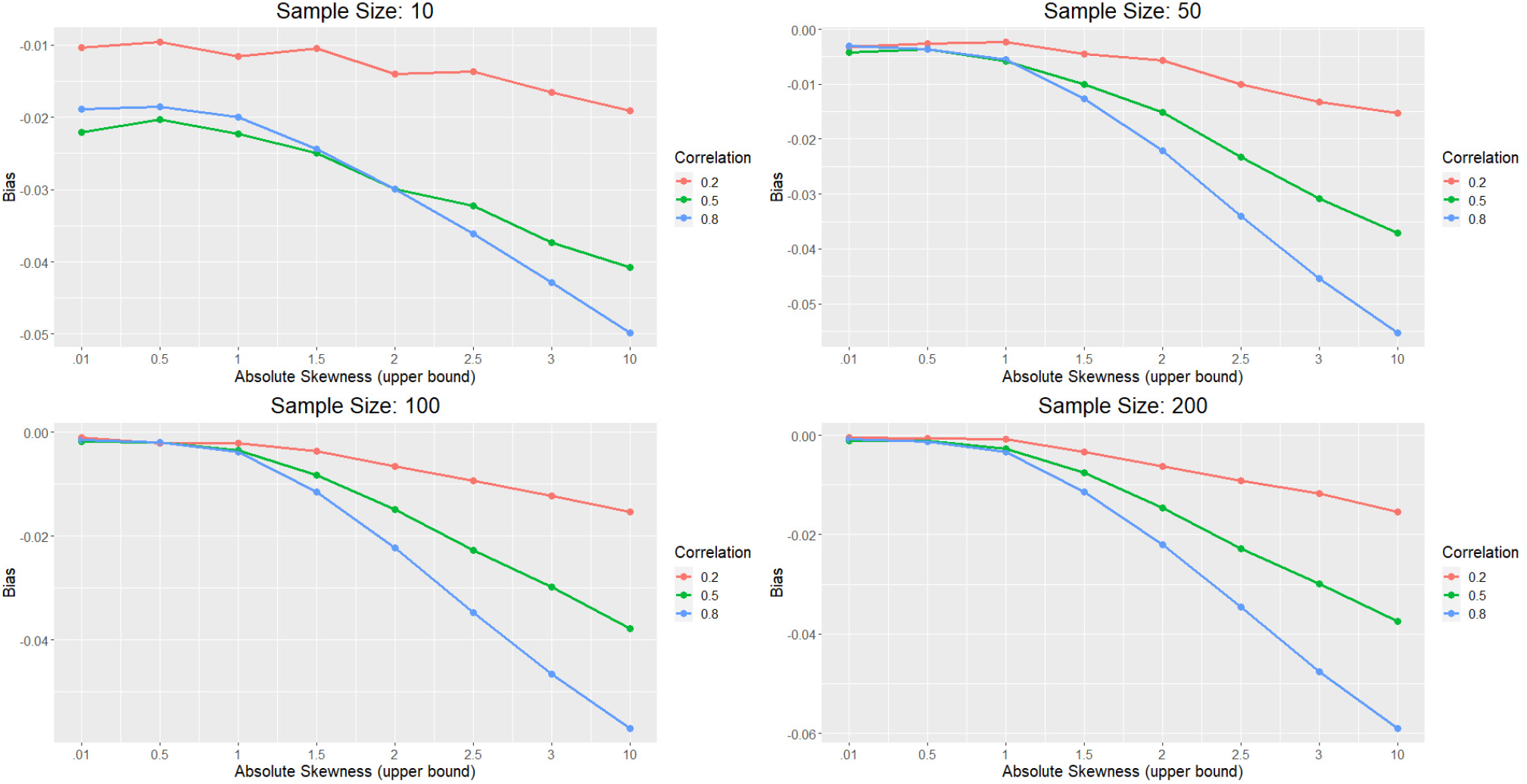
Correlation estimation results using the inverse CDF method. When skewness is low, there is very little bias in the correlation of the generated datasets. However as absolute skewness grows, so does the bias, getting as low as *−*.06 for the most extreme scenarios.

Figure 3 shows the results of this simulation. Unsurprisingly, the bias for each setting is always negative, confirming that this method tends to underestimate correlation. When the skewness of the beta distribution is low (*γ <* 1), the bias is very close to 0, particularly for large sample sizes. As the skewness increases, the bias becomes more severe, reaching as high as *−*2%, *−*4% and *−*6% for small, medium and large correlations, respectively. It should be noted, however, that absolute skewness levels above 2 or 3 are very rare in the reference data (0.61% and 0.14% frequency, respectively), while levels above 10 are nearly non-existent (*<* .01% frequency). Therefore, average bias on the whole tends to be quite small.

Absolute skewness levels of each dataset are reported in figure 1. While the median of each dataset is well below 1, certain datasets (lymphoma, newborn blood and sperm) have a small amount of CpG sites with relatively high (*>* 1) skewness levels. These distributions tend to generate slightly smaller linear correlations consistent with the results of the simulation, which in turn produce lower power levels. Given the rarity of these cases, the effect of these high-skewness distributions are marginal and tend to be washed out by the high frequency of distributions with low or moderate skewness.

### 3.2. Computation time

The second simulation compares the computation time required to calculate power across various sample sizes and settings. All settings assumed 100,000 CpG sites, with *K*_*m*_ *∈* (10, 100, 1000) of those having a non-zero correlation with the phenotype (i.e.,). Sample sizes of *n ∈* (10, 50, 100, 200) and correlations *ρ ∈* (0.1, 0.3, 0.5, 0.7, 0.9) were considered. The simulation also ran with a static *ρ* (*σ*_*ρ*_ = 0), and again with *σ*_*ρ*_ = 0.05. For all settings, an FDR of 5% was used with a target power margin of error of 1%. Because of this, each simulation setting used a different number of datasets (minimum is 20 and maximum is 1,000).

Partial results for the simulation are shown below in Table 2. Only simulation settings with 1,000 significantly associated CpG sites and with *σ*_*ρ*_ = 0 are shown (relaxing the latter restriction to *σ*_*ρ*_ = 0.05 increases the computation time by an average of 3 seconds). The number of datasets used in each setting is also provided in the parentheses, as this influences the overall computation time. Generally speaking, larger sample sizes and larger *K*_*m*_ correspond to longer run times. The same is not true for *K*_*n*_. In cases where the sample size is considerably small or large based on *ρ*, almost all datasets will lead to estimated power of 0 or 1, and the minimum number of datasets (*N* = 20) will typically suffice. This highlights another major advantage of EpipwR: the minimal cost of specifying a wide range of sample sizes, as the algorithm produces results from these ‘trivial’ power scenarios in just a few seconds. In cases where the average power is closer to 0.5, more datasets are needed to produce reliable estimates of average power, leading to longer computation times. Alternatively, increasing the target margin of errror will decrease the number of datasets required and therefore reduce the total computation time. At the recommended level of 3%, *N* is almost always under 300. Regardless of the scenario, the standard for EpipwR is to use anywhere from a few seconds to a few minutes to produce results, whereas other simulation-based tools can take hours or even days.

**Table 2.** Computation time in seconds of EpipwR for various settings with *K*_*m*_ = 1000.

| | $\rho = 0.1$ | $\rho = 0.3$ | $\rho = 0.5$ | $\rho = 0.7$ | $\rho = 0.9$ |
| --- | --- | --- | --- | --- | --- |
| $n = 10$ | 1.32 (N = 20) | 1.70 (26) | 28.39 (452) | 62.56 (1000) | 1.37 (20) |
| $n = 50$ | 4.64 (20) | 39.22 (172) | 87.37 (382) | 4.60 (20) | 4.66 (20) |
| $n = 100$ | 8.73 (20) | 170.20 (393) | 8.72 (20) | 8.67 (20) | 8.72 (20) |
| $n = 200$ | 17.49 (20) | 69.61 (82) | 17.07 (20) | 17.02 (20) | 17.09 (20) |

## 4. Discussion

EpipwR is the first open-source tool specifically designed for sample size determination of EWAS with continuous phenotypes. Similar to many power analysis tools for large-scale studies, it employs simulations to replicate empirical studies. But because these are limited to non-null and a small number of null tests, EpipwR is much faster than existing, fully-simulated EWAS power analysis tools (Graw et al., 2019). It is important to note that, like all power analysis tools, EpipwR makes several simplifying assumptions, and its outputs should be viewed as preliminary estimates rather than exact calculations.

Although EpipwR is introduced in this manuscript as a power calculator for continuous phenotypes, a future version of this package will contain tools to estimate power under a case control study design as well. Additionally, the methodology of EpipwR can be applied to much more sophisticated scenarios, such as the inclusion of one or more confounding variables, estimating correlation across CpG sites that can not be explained by the phenotype, and more complicated study designs (ANOVA, mixed effect models, etc.).

## Notes

### Competing Interest Statement

The authors have declared no competing interest.

